# FoldARE, an RNA secondary structure analysis and prediction tool via generative pseudo-SHAPE modeling

**DOI:** 10.64898/2026.03.04.709501

**Authors:** Stefano M. Marino, Vladyslav Husak, Toma Tebaldi

## Abstract

RNA secondary structure prediction is limited by conformational heterogeneity and the scarcity of experimental data, as many RNAs populate ensembles of near-isoenergetic folds and SHAPE data are often unavailable.

We present *FoldARE* (Folding and Analysis of RNA Ensembles), a two-step framework that derives pseudo-SHAPE constraints from in silico structural ensembles and uses them to guide SHAPE-aware secondary structure prediction. In the first step, an ensemble is generated and parsed nucleotide by nucleotide to estimate single-strandedness frequencies, which are converted into a pseudo-SHAPE reactivity profile using a weight-and-threshold scheme. In the second step, this profile is provided as a constraint to a SHAPE-compatible folding algorithm to improve the final prediction. We systematically evaluated all combinations of four ensemble-capable predictors, ViennaRNA, RNAstructure, LinearFold and EternaFold. After parameter optimization on a structurally diverse 25-RNA training set and validation using multiple scoring schemes, the best configuration combined EternaFold as ensembler and RNAstructure as predictor. Across external benchmark datasets (RNAstrand, ArchiveII and bpRNA) and the experimentally derived eFold dataset, *FoldARE* achieved the highest accuracy. Beyond prediction*, FoldARE* provides modules for ensemble-focused comparative analysis, including pairwise and multi-tool consensus assessment, per-nucleotide variability metrics, and interactive visualizations. Notably, it also supports the evaluation of m6A modification effects on structural ensembles.

*FoldARE* is freely available on GitHub (https://github.com/TebaldiLab/FoldARE) and as a web accessible version (https://rdds.it/foldare/)

## INTRODUCTION

The prediction of RNA three-dimensional structure is a challenging task, due to RNA inherent flexibility, with high rotational freedom across multiple dihedral angles, as well as a relatively (i.e. to proteins) limited structural diversity (a limited vocabulary of nucleotides, giving rise to a limited set of modular motifs) (Chen 2008; Ou et al. 2022); the latter feature, resolves into a smaller set of constraints to apply to the folding process, when computationally approximated. At the same time, RNA molecules show hierarchical folding: secondary structures (e.g. double strands and loops) form first and later assemble into a tertiary structure (Boerneke and Weeks 2018); this makes secondary structure predictions easier, as complicated three-dimensional arrangements can be neglected while assessing regions of complementarity (e.g. secondary structures, such as stems). Physical and chemical laws (e.g., for polar interactions, bulkiness) as well as empirically derived constraints or sequence co-occurrences have been employed to predict the formation of dsRNA portions, e.g. stems, or preferences for unpaired regions, e.g. loops (Mathews et al. 2004; Mathews and Turner 2006; Mathews et al. 2010). However, because of the aforementioned flexibility, the arrangement of secondary motifs in tertiary structures can take a multitude of alternative conformations, with comparable thermodynamics; the latter feature is a trademark of RNAs (Mustoe et al. 2014; Bonilla et al. 2024): in a given environment, RNA molecules can exist in different, almost isoenergetic, arrangements; an overall representation of these in structural models takes the name of a structural ensemble. A second aspect behind the challenges of RNA structural predictions is technological, in particular the significant limitation in the availability of experimental data, with only a fraction of the biomolecules solved in the Protein Data Bank (PDB) repository being RNAs (and in most cases, highly redundant for a few types, e.g. riboswitches, small stem-loops); this is due to intrinsic experimental difficulties (besides the high flexibility, other factors include heterogeneous ion binding networks, within an RNA population), for common structural biology techniques. Ultimately, the very low representation of unique structures for RNAs in the PDB, is still one of the biggest obstacles for current-day structural predictions (Jackson et al. 2023; Wang et al. 2023; Cruz et al. 2023; Schneider et al. 2024). Despite these challenges, predicting RNA structure is pivotal because an RNA’s function is tied to its form: RNA folding directs the function in fundamental biological processes (Cech and Steitz 2014; Bartel 2018; Statello et al. 2021; Morris and Mattick 2024); moreover, accurate structural models are a sought-after feature in drug discovery, to identify novel targets on RNA molecules, rather than or in alternative to protein targets (Warner et al. 2018; Childs-Disney et al. 2022; Disney 2025) or for RNA-therapeutics, where RNA is an active moiety itself (Baden et al. 2021; Haseltine et al. 2024).

Thus, the development of effective computational tools has been pursued since the 1970s with a focus on secondary structure predictions. Early methods relied on simple base-pairing rules (Watson–Crick), ignoring free-energy calculations (Nussinov Algorithm, 1978; Comay et al. 1984). A major advance came with thermodynamic modeling via dynamic programming, notably the Zuker Algorithm (Zuker and Stiegler 1981), which introduced free-energy minimization based on experimental parameters. Other strategies incorporated additional information, such as evolutionary signals from multiple sequence alignments in covariance models (Eddy and Durbin 1994; Nawrocki and Eddy 2013; Griffiths-Jones 2003). Machine-learning approaches followed, including CONTRAfold’s conditional log-linear models (Do et al. 2006) and, more recently, deep-learning tools such as SPOT-RNA and MxFold2 (Singh et al. 2019; Sato et al. 2021), with recent reviews summarizing advances in ML-based RNA structure prediction (Zhang et al. 2024; Wang et al. 2025). For energy-based methods, the two most established and actively developed packages that evolved from Zuker’s dynamic programming approach are ViennaRNA (RNAfold; Hofacker et al. 1994; Lorenz et al. 2011) and RNAstructure (Fold; Mathews et al. 1999). Both have added capabilities over time, including co-folding and consensus structure prediction from alignments. Most relevant for this study is their ability to integrate experimental data such as icSHAPE (in vivo click Selective 2’-Hydroxyl Acylation and Profiling Experiment) structural probing. In SHAPE experiments, a reactive probe (e.g., NAI-N3) electrophilically attacks the 2’-hydroxyl of RNA ribose residues. This nucleobase-agnostic reactivity correlates with nucleotide flexibility or accessibility, with high reactivity indicating single-stranded or flexible regions. Sequencing then identifies reactive sites as mutations or stops, allowing read coverage to infer per-position reactivity. This high-throughput approach probes RNA structure in cells, revealing flexibility, accessibility, and tendencies toward single-strandedness (Merino et al. 2005; Spitale et al. 2015). Such structural probing data can guide computational secondary structure predictions, a capability we specifically utilized in this study.

In omics-scale studies, high-throughput data at the bulk or single-cell level can capture transcript abundance and translational activity; however, once accumulating or depleting RNAs are defined, their functional and structural characterization is fundamental for a full biological understanding. Beyond mechanistic studies of individual transcripts, broader structural assessments (such as quantifying total double-stranded RNA accumulation across different cell types) allow for valuable cross-comparisons with expression data. While tertiary structure prediction remains unfeasible at this scale, many secondary structure tools are also unsuitable for omics applications due to software availability gaps, strict RNA length constraints, or the escalating computational burden that limits high-throughput workflows (as seen with tools like SPOT-RNA2, Singh et al. 2021).

The main objective of this work was to explore computational approaches suited for this task. In doing so, we designed and developed a novel, two-step predictive strategy that builds on existing methods by leveraging computational ensembles to generate pseudo-SHAPE data — reactivity tables that assign a score to each sequence position based on predicted reactivity. These pseudo-SHAPE profiles are then used as a co-input to guide a second, refined prediction step. The algorithm was first tested and parameterized using an in-house benchmark dataset, then evaluated on larger datasets, where it demonstrated improved performance compared to the individual baseline methods. We implemented the algorithm into a computational tool, called *FoldARE* (for Folding and Analysis of RNA Ensembles), designed for comprehensive secondary structure analysis. Beyond prediction (enabled by the newly developed *foldare_predict* algorithm). *FoldARE* supports customizable structural comparisons and statistical evaluation of alternative structural ensembles, including analysis of the structural effects of user-specified RNA modifications. The platform, available both as standalone and as web service, provides extensive output formats and interactive visualizations to enhance interpretation and facilitate comparison of predicted structures.

## RESULTS

We first introduce the two-step pseudo-SHAPE generative strategy underlying *FoldARE* predictions, then describe its parameterization and optimization, and finally assess its performance across multiple benchmark datasets and analysis tasks.

### Overview of the two-step pseudo-SHAPE generative strategy

The main idea behind our work was to evaluate the hypothesis that computational generation of RNA structural ensembles can be used to “mimic” SHAPE reactivity data, i.e. tables specifying positions and their relative tendency to be found in single-strand conformation. This can be achieved by building an *in silico facsimile* of a SHAPE experiment derived co-input: a text file with position related scoring, with higher scores for higher single strand frequencies (from SHAPE experiments, a high score describes a highly probe-reacting position); as previously introduced, similar co-inputs can then be included in secondary structure predictions, to guide them by “signaling” more probable single strand positions. In this work, we envisioned a two-step predictive approach (**Figure 1**) where, in the first step, ensemble data are generated, and then used to create a table reporting positions with “single-strandness” tendencies, i.e. positions (more or less) likely to stay single-stranded throughout different conformations, within a predicted ensemble. These *in silico* estimated tendencies are saved in a text file, working as a *pseudo* SHAPE co-input (i.e. a co-input guide file) for a second run of prediction, by employing currently available SHAPE-aware predictive algorithms. The methods considered were chosen as they could comply with these requirements: the ability to generate structural ensembles (i.e. a set of alternative conformational models, explicitly laid out) as well as the capability to accept SHAPE-derived data in the form of text-based co-input files described before, to ultimately generate SHAPE-aware predictions. The methods fitting these criteria (more details in the Methods section) were: ViennaRNA (RNAfold and RNAsubopt modules), RNAstructure (Fold module), EternaFold (Wayment-Steele et al. 2022), LinearFold (Huang et al. 2019). We designed our strategy by combining the four methods in “ensembler-predictor” pairs (overall 16 combinations; **Figure 1**), where “ensembler” is the method generating the ensemble (to be parsed for creating the pseudo-SHAPE co-input), and “predictor” is the method used to take in the pseudo-SHAPE co-input to assist the prediction. We developed the algorithm in parametric form, where each combination was tested against a test dataset (described in the next paragraph) with variable weights and cut-offs. As shown in **Figure 1**, the steps of our approach are: *(i)* a method (“ensembler”) is run in “ensemble mode”, i.e. returning an ensemble in output; then *(ii)* the ensemble of alternative models is parsed for each position to determine the frequency of single strand (ssRNA) predictions, e.g. relative frequency of “.” symbol, within the ensemble.

**Figure 1.**
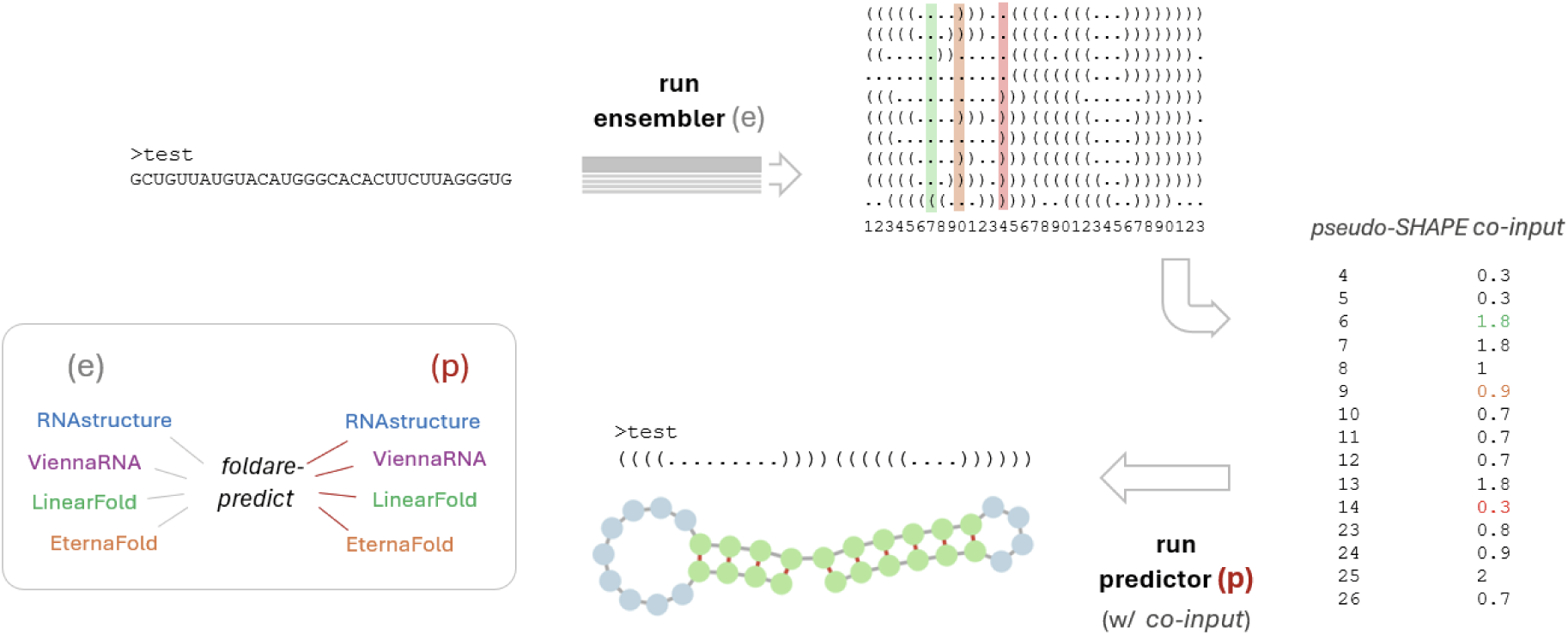
Overview of the two-step pseudo-SHAPE generative strategy. Given an input sequence, the first step is to run an ensemble generating method (“ensembler”, (e)); the generated ensemble is parsed, position per position (indexes are shown below the ensemble, 0 to 9, for each set of ten), evaluating frequency, and through this compute a pseudo-SHAPE reactivity value, reflecting the proportion of ssRNA in the ensemble (see color coded examples in the figure); positions for which the ssRNA frequencies are low (< chosen cut-off) are skipped, the resulting values are stored in a text file (the pseudo-SHAPE co-input) then used for a second step: the co-input is taken in by a predictive algorithm (“predictor”, (p)) as a guide for new predictions. The 4 selected methods are reported in the box at the bottom left of the figure; they can be combined in 4×4 combinations, yielding 16 different alternative predictive structures.

### Parameterization and optimization of the *FoldARE* predictive framework

Parametric cut-offs (x1, x2, x3) and weights (k1, k2, k3) were set to determine the limits of f(ssRNA) to consider and their relative impact on the pseudo-SHAPE score. For example, a scheme: x1=0.9, x2= 0.5, x3=0.3, k1=2, k2=1, k3=0.5, implies, for each positions, that within-ensemble frequencies of ssRNA predictions f(ssRNA) above 90%, 50%, 30% are given a relative weight of 2.0, 1.0 and 0.5 respectively (thus a frequency of single strand < 30%, is skipped). This scheme, based on an “educated guess”, was tested first as a proof of concept (see later on, discussion on **Figure 2A**). During the “ensembler step”, relative weights and cut-offs are applied to each position, evaluating each f(ssRNA)_i_: this creates positional pseudo-SHAPE scores, then reported in the pseudo-SHAPE co-input file (see example in **Figure 1**; further details in the Methods section). In essence, this co-input reports all positions with an above cut-off degree (set by the x3 parameter, in the previous example, x3=0.3) of single-strandedness, with higher values for higher frequencies (and null values for any position with f(ssRNA) < x3). Ultimately, the pseudo-SHAPE co-input assists the final prediction, which returns the best (MFE or MEA) structure in dot-bracket notation. The approach defines three bins: high f(ssRNA) values (> x1), to which to apply the highest weight (k1); middle range f(ssRNA) values (from x1 to x2), to which to apply a middle weight (k2); lower range f(ssRNA) values (from x3 to x2), to which to apply the lowest weight (k3).

**Figure 2.**
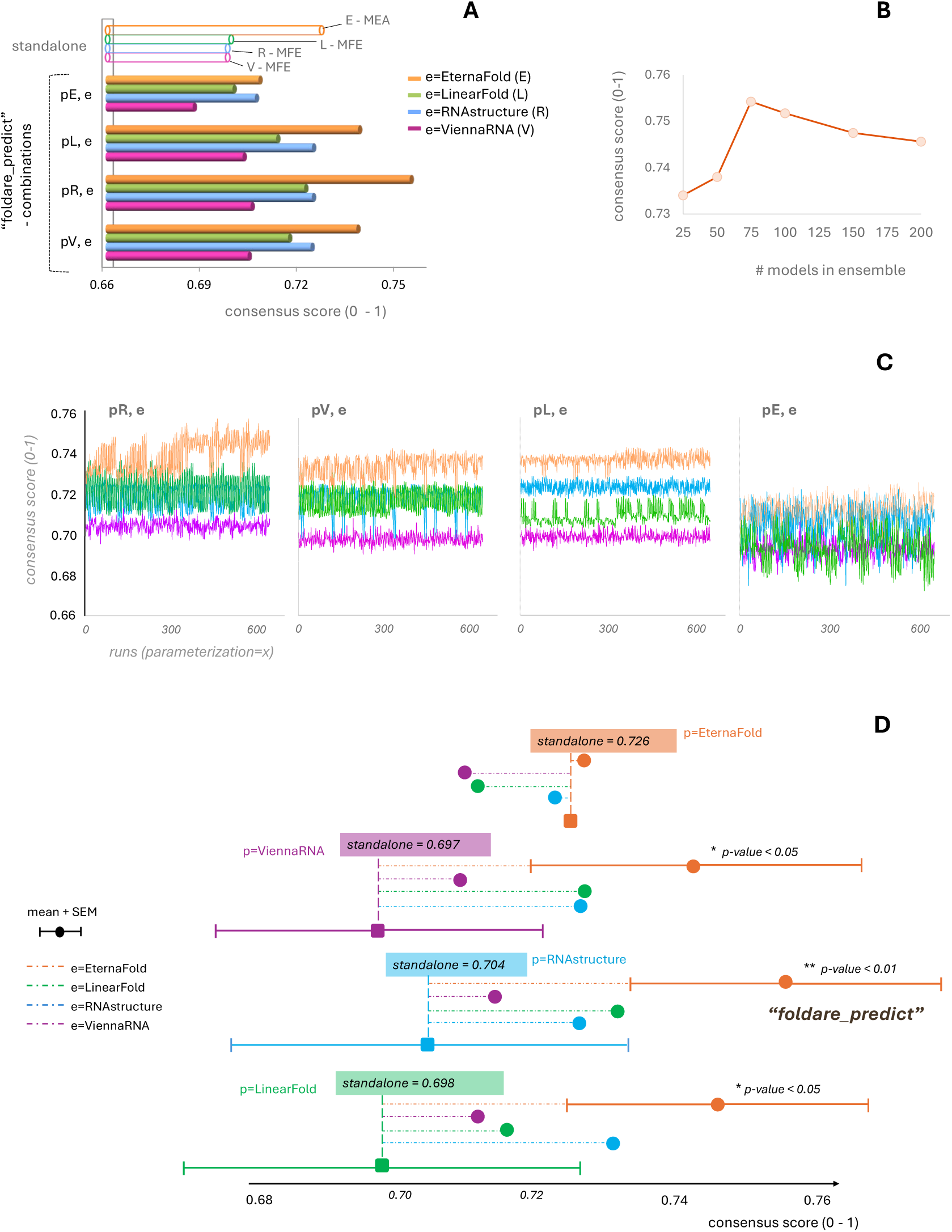
Parameter optimization of the ensemble-derived pseudo-SHAPE framework. **A**: the consensus scores (y-axis, 0 to 1, fraction of correct predictions, mean value for the 25db) is reported, for standalone methods (top of the figure; MEA and MFE: maximum expected accuracy and minimum free energy, for best models produced) and each of the 16 (predictor x ensembler) combinations (p=predictor, e =ensembler), with the initial (“educated guess”, pre-optimization) parameterization. **B**: effect of the ensemble size, for the top scoring combination: p=RNAstructure, e=EternaFold (foldare_predict). **C**: optimization of parameters (cut offs: x1, x2, x3; and weights: K1, K2, K3; ensemble size: 50, 75) for each of the combinations: in the x-axis each point represents a run, with a unique parametric scheme; the red arrow marks the significant step, from ensemble size of 50 to 75 (rightmost half of the runs), for the, best scoring combination. In this panel, one graph for each predictor (abbreviations and color coding as in the legend of panel A) is reported (in each graph: four runs for any of the four ensemblers). **D**: comparative evaluation (by consensus score) of all 16 combinations of predictors/ensemblers with optimized parameters (as from **C**); mean and SEM are reported for standalone predictions (vertical dashed bars) and for best ensembler/predictor combinations (for each predictor); statistically significant (t-test) results are marked (*)

To develop and tune the method, we needed to define a reference dataset on which to run the algorithm in its parametric form and to define the most effective scheme for cut-offs and weights. Thus, a manually curated database of 25 RNA structures was assembled by selecting entries from RNAstrand (containing SRPDB and Rfam data; details in Methods section) and the PDB; to ensure structural diversity and minimize redundancy, we applied the following selection criteria: sequence identity < 60%, 1500 nt > length > 50 nt, and a double-stranded content of at least 20%. From the resulting pool, we manually selected 25 models to capture a broad range of molecule types and biological sources, intentionally avoiding over-representation of any single RNA family (**Table 1**; detailing the “25db” dataset), an important consideration given the excessive redundancy of some types. The limited dataset size was chosen to keep the downstream optimization runs within a reasonable computational time.

**Table 1.** 25RNA structures (25db) for the development and optimization of the algorithm.

| Name | Source | Length (nt) | Description |
| --- | --- | --- | --- |
| ASE_00001.dp | RNA STRAND | 262 | RNase P RNA, *Acidianus ambivalens*, strain DSM 3772 |
| ASE_00251.dp | RNA STRAND | 189 | RNase P RNA, *Pichia guilliermondii*, strain NRRL Y-2075 |
| CRW_00212.dp | RNA STRAND | 1528 | Gutell Lab CRW, file name: d.16.b. S.griseus.bpseq, ID: X61478 (5S rRNA, *S. griseus*) |
| CRW_00438.dp | *RNA STRAND* | 954 | Gutell Lab CRW, d.16.m.<br>H.sapiens.5.bpseq (mitochondrial tRNA, *H. sapiens*) |
| PDB_00897.dp | *RNA STRAND* | 152 | ribosomal protein L1 *Thermus thermophilus* |
| PDB_00904.dp | *RNA STRAND* | 152 | P-site and P/E-site tRNA structures |
| PDB_00908.dp | *RNA STRAND* | 222 | group I intron/two exon complex (human) |
| PDB_01039.dp | *RNA STRAND* | 1511 | 30S ribosome |
| PDB_01112.dp | *RNA STRAND* | 75 | riboswitch (E.coli) |
| PDB_01118.dp | *RNA STRAND* | 57 | H/ACA ribonucleoprotein particles (RNPs) |
| SRP_00030.dp | *RNA STRAND* | 272 | *Paenibacillus macerans* scr gene for small cytoplasmic RNA |
| SRP_00035.dp | *RNA STRAND* | 267 | *Bacillus stearothermophilus* small cytoplasmic RNA |
| SRP_00188.dp | *RNA STRAND* | 250 | *Lycopersicon esculentum* mRNA for signal recognition particle 7S RNA (variant L) |
| SRP_00199.dp | *RNA STRAND* | 320 | TPA: *Methanococcoides burtonii* DSM 6242 SRP RNA for signal recognition particle RNA |
| TMR_00192.dp | *RNA STRAND* | 355 | TPA: *Shewanella* sp. ANA-3 tmRNA gene Shewa_sp_ANA3 |
| 1NBS | *PDB* | 155 | *Bacillus subtilis* Ribonuclease P RNA |
| 1VTQ | *PDB* | 75 | Yeast (*Saccharomyces cerevisiae*) tRNA-ASP. I. |
| 3DIG | *PDB* | 174 | *Thermatoga maritima* Lysine Riboswitch |
| 3IWN | *PDB* | 93 | Co-crystal structure of a bacterial (*Vibrio cholera*) c-di-GMP riboswitch |
| 3PDR | *PDB* | 161 | M-box RNA (*B. subtilis*) |
| 4KQY | *PDB* | 119 | yitJ S box/SAM-I riboswitch (*B. subtilis*) |
| 7SHX | *PDB* | 94 | *H. sapiens* sense-oriented promoter-associated non-coding RNA (paRNA) |
| 7ZFW | *PDB* | 616 | *H. sapiens* 80S ribosome |
| 8TJV | *PDB* | 417 | *Thermotoga petrophila* Ribozyme scaffolded Fluoride riboswitch (2023) |
| 9CBU | *PDB* | 387 | *Tetrahymena thermophila* ribozyme (2024) |

We tested the 2-step algorithm in its parametric form, code named “foldare_predict_XY” (where foldare_predict is our predictive algorithm, and X and Y stand for the different combinations of X=“predictors” and Y=“ensemblers” employed), with previously defined cut-offs ( x1, x2, x3) and weights ( k1, k2, k3) set to vary in the ranges (0.9, 0.8, 0.7), (0.6,0.5), (0.4, 0.3, 0.2), (3, 2.5, 2),(1.75, 1.5, 1.25), (0.75, 0.5), respectively. Additionally, the ensemble size was set to (50, 75). For each run, i.e. for each parametric combination and for each “foldare_predict_XY” combination, the predictions were compared against the ground truth, i.e. known reference structures in the 25db. In particular: for each 25db entry, comparing the predicted secondary structure to the known reference, identical predictions were counted (+1, for correctly assigned dsRNA opening positions “(“, or closing positions “)”, or ssRNA, “.”), and normalized for the total length of the sequence, ultimately reflecting the fraction (0 to 1) of correct predictions, for each entry in the 25db. We termed this scoring system, the simplest and fastest scheme, consensus score. With it, we evaluated all the runs of the parametric optimization of “foldare_predict_XY”; results of these optimizations are shown in **Figure 2**; panel A (with full details reported as Supporting Information, Table S1) reports the best scoring combination, for each (4×4) combination of “ensembler” and “predictor” in respect to the 25db, as well as the results for each predictor run as a standalone program (with default parameters, as by its most common employment). As for the latter: the most similar predictions (evaluated by correlating the predicted scores for each entry in 25db), were those from ViennaRNA (RNAfold) and RNAstructure (Fold), with a Pearson Correlation Coefficient (PCC) =0.936; this is not a surprise, considering the similarity between these two (dynamic programming, thermodynamic-based); then, LinearFold with ViennaRNA, PCC=0.821 and LinearFold with RNAstructure, PCC=0.788; the most diverging method was EternaFold (with RNAstructure, PCC=0.722; with ViennaRNA, PCC=0.698; with LinearFold, PCC= 0.654). Furthermore, considering the accuracy by consensus scoring: LinearFold, RNAstructure, and ViennaRNA returned very close performances (on average a score of 0.70, meaning that ∼ 7 out of 10 positions were predicted correctly, as compared to the reference in the 25db), while EternaFold returned the best performance (0.725). Aside from the different derivations (thermodynamic-based, machine-learning-assisted, etc.), results were close, indicating that these methods yield similar performance, regardless of their theoretical background.

Focusing on the foldare_predict_XY framework, Figure 2A displays the scores for the various combinations derived from the initial “educated guess” parametric scheme. In all instances, the highest performance was achieved using EternaFold as an ensembler. These results were obtained using an ensemble size of 75, a value that was identified as optimal in preliminary analyses and confirmed as the most effective for the 25db dataset (Fig. 2B); this parameter determines the number of alternative models considered for generating the pseudo-SHAPE data, prioritizing thermodynamically favorable structures (either via explicit free-energy ranking or stochastic sampling, see Methods). While an ensemble that is too small limits the representation of structural diversity, an excessively large ensemble introduces noise from lower-ranked models. Figure 2C displays the results of the multiparametric optimization, where cut-offs and weights were varied across defined ranges ( x1: {0.7, 0.8, 0.9}, x2: {0.5, 0.6}, x3 :{0.2,0.3,0.4}, K1:{2, 2.5, 3}, K2: {1,1.25}, K3: {0.5, 0.75}, ensemble size: {50, 75}).

Full details are provided as Supporting Information, Table S2. The optimization results indicate two key trends: *(i)* the top-scoring combinations invariably feature EternaFold as the ensembler (indicated by the orange line in Fig. 2C, which outperforms all other combinations: for RNAstructure, for ViennaRNA, for LinearFold, and for EternaFold); and *(ii)* RNAstructure is the most parameter-sensitive predictor, with a distinct preference for an ensemble size of 75, confirming previous results (Fig. 2B) Taken together, these runs established the most effective parameterization for our algorithm - hereafter referred to as *foldare_predict* (specifically the p=RNAstructure, e=EternaFold configuration). The optimized parametric scheme (x1=0.9, x2=0.5, x3=0.3, K1=2.5, K2=1.25, K3=0.75) closely aligns with our original “educated guess.”

Notably, the algorithm’s design (specifically the pseudo-SHAPE co-input) effectively “steers” the calculations, with (step 1) co-inputs significantly assisting (step 2) predictions. Correlations between *foldare_predict* and standalone versions of EternaFold (PCC = 0.931) and RNAstructure (PCC = 0.821) demonstrate that the ensembler-based co-input yields results distinct from either program alone. Beyond the primary *foldare_predict* configuration, other combinations also outperformed standalone predictors. As shown in Figure 2D, the mean and standard error for the 25db dataset reveal significant improvements: while *foldare_predict* achieved the highest significance (p < 0.01), both ViennaRNA and RNAstructure benefited significantly from EternaFold integration (p < 0.05). These findings underscore that a two-step consensus approach effectively smooths out dissonant predictions and reinforces structural accuracy against the ground truth.

### Robustness to scoring schemes and evaluation metrics

At this stage, we aimed to further refine the analysis: in particular, to strengthen the validation, we examined and quantified the effect of the chosen scoring system on the results, an important aspect to consider. We decided to perform a second run of parametric optimization, this time refining the search to the top-performing *foldare_predict* configuration by employing closer intervals (of the xs and Ks parameters) around the best-performing values in Figure 2C, while assessing the results in parallel with alternative scoring systems. Specifically, *(i)* we run the optimization by varying each parameters in the following ranges: x1 ={0.8, 0.85, 0.9}, x2 ={0.5, 0.55, 0.6}, x3 ={0.3, 0.35, 0.4}, K1 ={1.75, 2, 2.25}, K2 = {1, 1.25}, K3 = {0.5, 0.75}, all with ensemble size =75; and *(ii)* employed three different scoring methods; as for the latter, together with the consensus scoring previously described (i.e. an identity score, returning the % of identical prediction; here, in short scoreA), we employed a similarity score (scoreC), assessing structural similarity within a 5-nucleotide window, accommodating minor shifts (+1,-1) in base pairing, and finally a matching bracket scheme (scoreD; a stricter form of identity score, requiring a bracket, representing a base pair, not only to be predicted correctly but also to be paired with the correct partner), which allows for the calculation of a standard F1-score (details and formulation in Methods). With these different schemes, we assessed the optimization runs (**Figure 3 A-C**; and Table S2); two very similar weight schemes gave the best results for all scoring systems: x2=0.5, x3=0.3, K1=2.25, K2=1.25, K3=0.75 for all, with x1=0.9 (best for scoreC, equally best for scoreA) or x1=0.85 (best for scoreD, equally best for scoreA). Comparative results against the 25db dataset of *foldare_predict* (with the optimized scheme above, x1=0.85) versus each of the standalone methods are reported in **Figure 3D**. Here, positive Z-scores represent performances that are better than average (the higher the Z-score, the better the performance), and conversely, negative Z-scores (i.e. reporting performances below average); overall, *foldare_predict* returned the best results by all the alternative scoring methods employed (**Figure 3D**).

**Figure 3.**
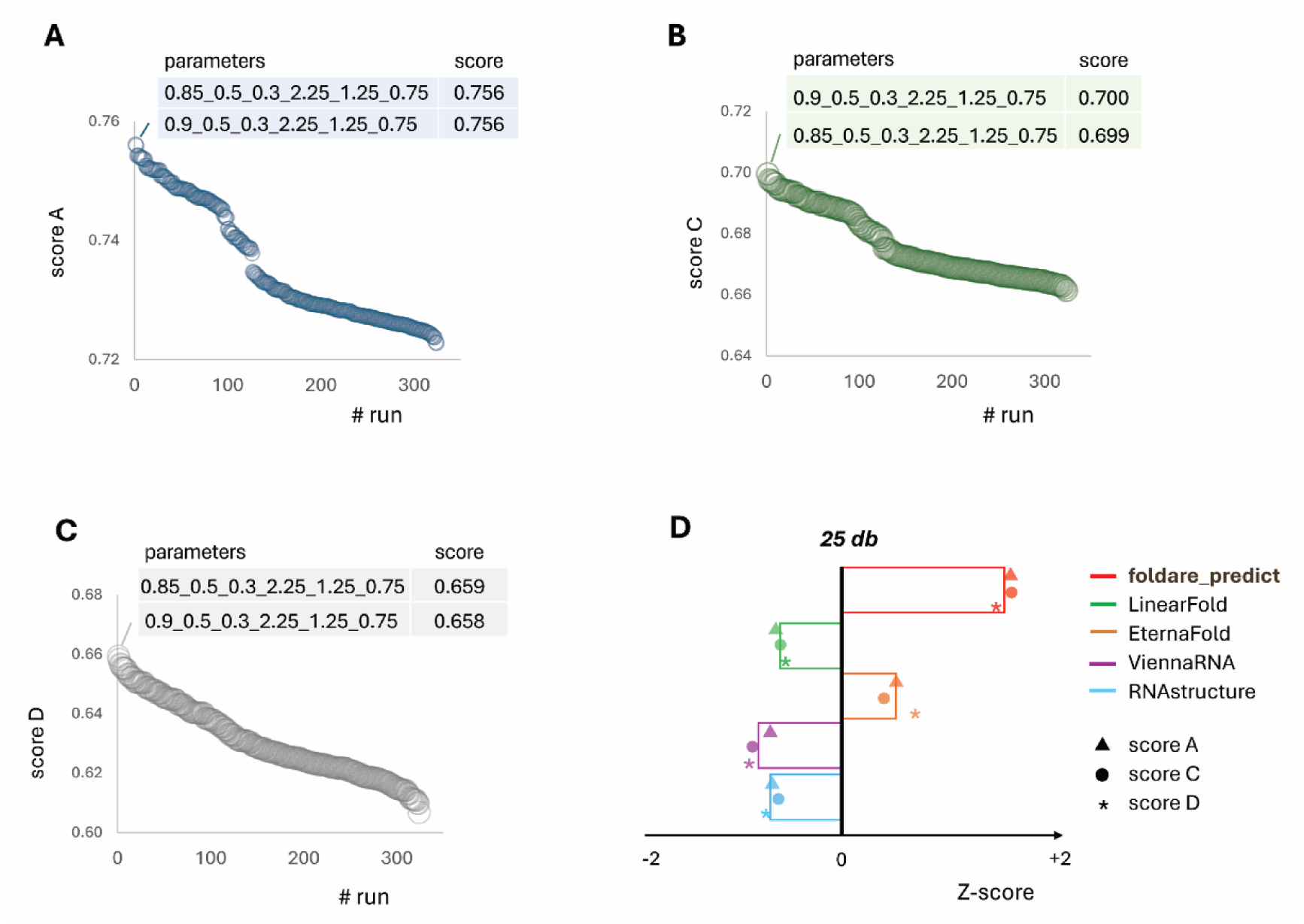
Robustness of FoldARE performance across scoring schemes. **A-C**: Validation of the approach applying different scoring schemes: identity score (% of identical predictions) in **A**; similarity score, reflecting local (5 nt long) similarity in **B**, and a matching bracket identity score, reflecting identity in terms of exact base-pairing, in **C**. Details on the scoring systems are provided in the Methods section. On the horizontal axis, simulation runs are reported (ordered by score, from high to low). On top of each plot, the highest-performing schemes are reported (x1_x2_x3_K1_K2_K3 for cut-offs and weights). **D**: performance evaluation with the 25db; for each predictor, bars represent the average Z-score for the 3 different scoring methods (the 3 symbols, in each bar); Z-score > 0 reflects better than average results.

### Benchmarking against established RNA secondary structure predictors

Next, with the refined scheme (x1=0.85, K1=2.25 in Fig. 3D), we moved on to a larger assessment with more extensive benchmarking. We employed different standard datasets of RNA secondary structures, taken from the literature and previously employed as reference datasets for similar predictions. These datasets were: RNAstrand (Andronescu et al. 2008), Archivell (Sloma and Mathews 2016), bpRNA (Danaee et al. 2018), all filtered for: redundancy (< 80% sequence identity), size selected, and with at least 20% content in dsRNA (for more details, see Methods). Additionally, we employed a more recent dataset, called eFold, including experimentally derived structural data (determined by chemical probing) for mRNA data for *H. sapiens*, pre-miRNAs and lncRNA (https://github.com/rouskinlab/eFold); thus, this data offers a unique opportunity to test available structural prediction methods on less-characterized human RNAs. Details of these datasets are provided in the Supporting Information, Table S3. Results are reported in **Figure 4A** (for the “standard” benchmark) and **Figure 4B** (for eFold): in all cases, *foldare_predict* topped the list of performance, with other methods (in particular, EternaFold and LinearFold), trading positions, depending on the case. Altogether considered (combining the benchmark datasets, n=2988), the results of *foldare_predict* were the best both in terms of improved accuracy (best Z-score, **Figure 4C**; datapoints distribution, in **Figure 4D**) and in terms of significance: as for the latter, *foldare_predict* returned the most significant performance increase in respect to the other methods, singularly considered, with p-value < 1*10^-50^ in all cases (p=2.7 e^-68^ with EternaFold, p=3.1 e^-52^ with RNAstructure, p= 1.6 e^-50^ with LinearFold, p= 1.9 e^-64^ with ViennaRNA).

**Figure 4.**
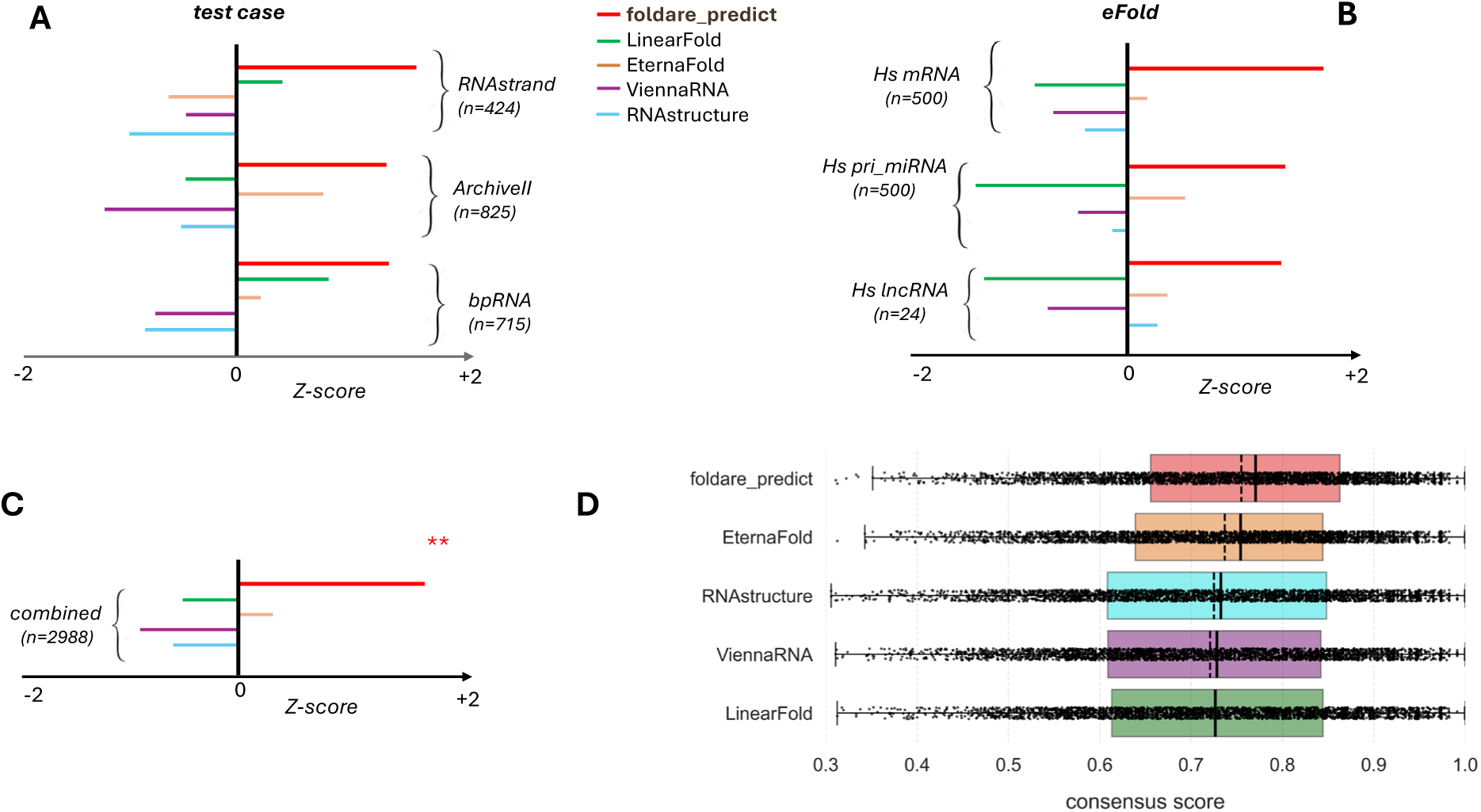
Benchmarking FoldARE against established RNA secondary structure predictors. Performance evaluation (for each predictor bars represent the average Z-score for the 4 different scoring methods tested, i.e. the four symbols, in each bar) with external reference datasets: (i) ArchiveII, RNAstrand, bpRNA in **A**, and (ii) eFold (human mRNA data, miRNA and lncRNA data) in **B**. The combined results (collecting all datasets in **A** and **B**) are reported in **C**, and were tested for significance in pairwise comparisons of values (Wilcoxon signed rank, one-tailed, testing the improved performance of foldare_predict vs any other method): in **C**, two stars flag a p-value < 1*10^-10^ for foldare_predict, in all comparisons. The legend for symbols and colors is reported in the figure. **D** shows the actual distribution of data points for the combined analysis, with average (dashed line) and median (line).

We note that the eFold dataset was not used during the development of any of the standalone methods, thus making it a form of blind test. We therefore used this opportunity to extend the benchmark to a state-of-the-art machine learning method, Mxfold2 (whose training set included the same datasets we employed in Fig 4A, but none from eFold, which was published subsequently), as a representative of recent deep learning algorithms in RNA secondary structure prediction (Sato et al. 2021). Notably, Mxfold2 is a pseudoknot-free method that is well-suited for large-scale analyses (it is free as a standalone tool, resource-efficient, and handles any RNA size). Against eFold, Mxfold2 achieved an average consensus score of 0.680 across the three datasets (Fig 4B; *n* = 1024), comparable to RNAstructure (0.692) and ViennaRNA (0.678), higher than LinearFold (0.664), but lower than EternaFold (0.702) and foldare-predict (0.731).

We also tested another common deep learning method, SPOT-RNA (Singh et al. 2019), for which the eFold dataset was not included in its training. SPOT-RNA returned an average consensus scores of 0.587, across the three eFold datasets, the lowest in the benchmark. It should be noted that SPOT-RNA is a pseudoknot aware method, meant to provide more “refined” structural descriptions (intermediate between secondary and tertiary structures), an important plus of this tool, that however went unused in this analysis (with the pseudoknot-free eFold reference); a major reason for this result, resides in that SPOT-RNA predicted a disproportionately high percentage of unpaired positions: in the combined eFold dataset it predicted an average of 70.3% single stranded positions, compared to the 44.3% of the ground truth, while all other methods returned much closer evaluations (40.3% mxfold, 44.8% foldare-predict, 46.5% EternaFold, 36.1% ViennaRNA, 49.2% LinearFold, 37.1% RNAstructure). Detailed results of these calculations on the eFold “blind test”, are provided in Table S4.

The *foldare_predict* algorithm is implemented in *FoldARE (*https://github.com/TebaldiLab/FoldARE), further described in the next section.

### Ensemble comparison and m6A-aware analysis with *FoldARE*

We developed the FoldARE tool *(i)* to provide predictive capabilities, and *(ii)* to compare and interactively visualize the output of the different established approaches considered in this study. The first capability is achieved by implementing the optimized algorithm described in the previous paragraphs (*foldare_predict*, **Figure 5**). *foldare-predict* is available as a command-line tool and, by default, implements the optimized configuration described above. However, its two-step prediction workflow is highly customizable through a configuration file and a set of optional parameters, allowing users to explore and test any of the combinations described in Fig. 2. In addition, we provide a web-accessible version that does not require computational expertise. Notably, alongside *foldare-predict*, the web platform includes *psSHAPEr*, a utility for custom pseudo-SHAPE data generation. Using this tool, users can upload a custom ensemble either in dot-bracket notation or as a base-pairing probability matrix; *psSHAPEr* then automatically computes matrix-derived pseudo-SHAPE profiles, generating a text-based input file that can be directly used by *foldare-predict*. The second capability (“foldare compare”) resides in a set of scripts for comparative analysis of RNA secondary structures; there are two modalities, one comparing standalone methods in pairs (“compare-pair”, **Figure 5**), and the other comparing all methods simultaneously (“compare_all”, **Figure 5**). In brief, “compare-pair” compares pairs of (user selected) predictive methods, chosen among the four available options (EternaFold, LinearFold, RNAstructure, ViennaRNA); the analysis returns different graphical outputs, including an interactive similarity heatmap (where all pairwise cases can be singularly investigated by hovering over with the mouse), as well as per-nucleotide consensus scoring, Shannon entropy, and single-strandness frequencies distribution (f(ssRNA), **Figure 5**).

**Figure 5.**
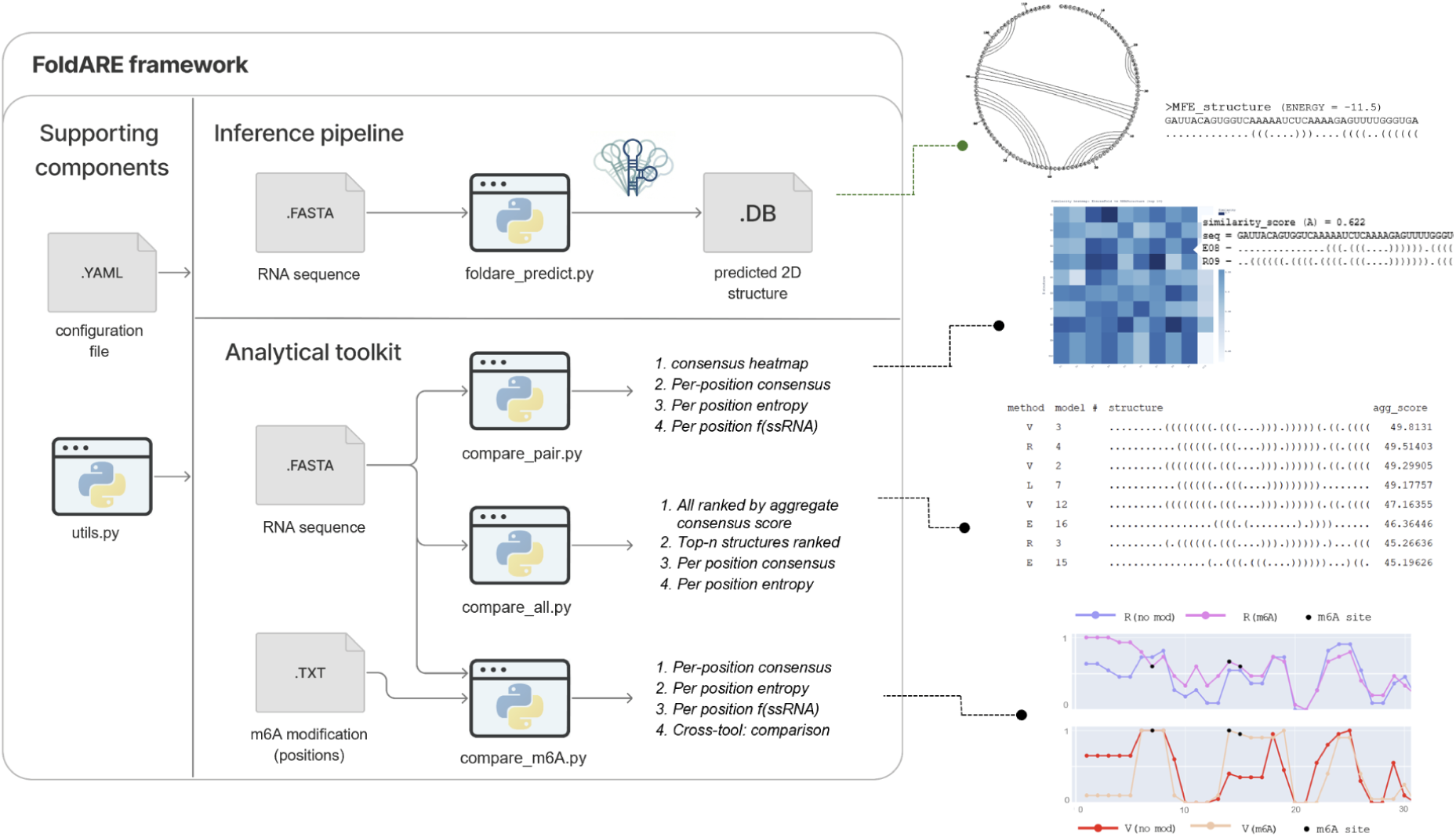
FoldARE enables ensemble-level comparison and modification-aware analysis. The main sections of the FoldARE computational tool described in the text: the framework includes four “modules”, one for prediction (inference pipeline, top), three for comparative analysis (Analytical toolkit, bottom section); output types are reported in the figure, with dedicated highlights (one for each “module”) on some of the most relevant output information (text and plots) described in the main text.

The “compare-all” mode compares the ensembles generated by all tools run in parallel. From their output, it computes the similarity score between any pairwise top hits, returning *(i)* a graphical and interactive representation of the average consensus per position across all different predictors, and *(ii)* an aggregate score as an output text file (Figure 4C), ranked from best to worst, with the highest-consensus models topping the list. Importantly, these analyses can be extensively customized. Users can choose different predictor combinations or ensemble sizes, as well as apply different scoring schemes to obtain the most robust results possible. While the consensus score (scoreA) is the default option, users can also opt for the similarity score (scoreC) or the matching bracket scheme (scoreD), for any of the comparative tools in *FoldARE*.

Last but not least, an additional feature of *FoldARE* is represented by a third modality, dedicated to RNA modifications (“compare_m6A”, in **Figure 5**); currently, this analysis can be done only for the most common RNA modification, m6A, and only through ViennaRNA and RNAstructure (currently, the only methods capable of handling it); with compare_m6A, given a sequence in input and a co-input text file listing the modification sites, *FoldARE* automatically builds the models (unmodified and modified) and then comparatively parses the generated ensembles; after that, it reports both tool-specific and cross-tool plots showing the effects of m6A modifications, with graphical and interactive outputs analogous to those in “compar-pair”, but specifically highlighting the positions with the modification (**Figure 5**). To our knowledge, *FoldARE* is currently the only available tool for structural comparisons in RNA modification studies at the ensemble level, and it also supports cross-tool evaluations.

Alongside the standalone tool, we developed a web-accessible version (https://rdds.it/foldare/) to facilitate use by researchers without a computational background. As previously introduced, the web interface provides full access to the predictive and analytical tools shown in Figure 5, while introducing a dedicated utility—called *psSHAPEr*—for generating custom pseudo-SHAPE co-inputs. This allows users to perform highly specific, customized investigations through a simplified browser interface, bypassing the need for local installation or command-line expertise.

## DISCUSSION

In this work, we evaluated existing predictive methods for their ability to assess RNA secondary structure not only as single favored conformations (e.g., MFE or MEA), but more comprehensively as structural ensembles.

We hypothesized that mimicking experimental methods for structural probing, such as those based on SHAPE reactivities, we could develop a computational approach to generate *(i)* an *in silico* pseudo-SHAPE input (with information derived from a first modelling run), then *(ii)* use it as a guide for a second run of modelling. In the first part of this work, we tested the idea that a two-step, pseudo-SHAPE generative approach could improve the accuracy of secondary structure prediction. We selected methods that meet essential requirements for the task, specifically the ability to output structural ensembles with explicit representations of each alternative structure, and support for SHAPE co-input files — text files providing per-position reactivity scores; the list of methods fitting these criteria included two thermodynamics-based approaches (ViennaRNA, RNAstructure) and two machine learning-based tools (LinearFold, EternaFold).

We first tested alternative designs, sampling all possible combinations of the methods as “ensemblers” and “predictors” with different weight schemes. The most effective, i.e. best in deriving pseudo-SHAPE data, ensembler was EternaFold, while the best, i.e. best in using the pseudo-SHAPE data as co-input, predictor was RNAstructure. This combination was optimized against a training dataset (the 25db) and became the core of our predictive strategy, called *foldare_predict*. This algorithm was then evaluated on larger reference datasets not used during training but commonly employed in the literature — ArchiveII, RNAstrand, and bpRNA — where it demonstrated improved performance over standalone predictors. Furthermore, we tested our method on a recent dataset of experimentally validated secondary structures (eFold), which includes SHAPE-mapped human RNAs (mRNA, lncRNA, miRNA). Also in this challenging benchmark, the two-step approach achieved the best results. We note that these improvements are, in some cases, marginal, with all methods returning close and comparably good performances (whether ML or thermodynamics-based). For example, in the combined analysis of all datasets in the “standard” benchmarking in **Figure 4A**, the consensus scores (scoreA) ranges from 0.734 (73.4% of consensus positions, for ViennaRNA) to 0.760 (76%) for *foldare_predict*, with other performances in between (74.7%, 74.8%, 73.6% for EternaFold, LinearFold, RNAstructure respectively); on the eFold dataset, the difference was more pronounced, with our two-step method achieving the highest accuracy (on average, across the three datasets, 73.1% for *foldare_predict*), outperforming EternaFold (70.2%), RNAstructure (69.2%), ViennaRNA (67.8 %), and LinearFold (66.4%).

The convergence in performance may stem from the fact that RNA secondary structure predictors are often trained and evaluated on similar standard benchmarks (e.g., ArchiveII, RNAstrand). Additionally, inherent limitations in dot-bracket notation, a simplified 2D representation of RNA structure, may constrain the diversity of predicted structural descriptions, potentially compressing the space for divergent or alternative conformations. Nevertheless, *foldare_predict* consistently outperformed across all conditions and cases; notably, the eFold dataset was not used during the development of any of the standalone methods, making it a good blind test. We took advantage of this to extend our analysis to two deep learning methods that were not trained on it (Mxfold2 and SPOT-RNA) ensuring a fair comparison across different approaches. In this eFold blind test, Mxfold2 achieved a fairly good performance (an average consensus score of 68.0%, comparable to the values previously outlined for ViennaRNA and RNAstructure), while SPOT-RNA returned a below-average score (58.7%). A thorough benchmarking of machine learning methods was beyond the scope of our work (which focuses on SHAPE aware, ensemble generating methods); nevertheless, these results would support the view that while deep learning methods often excel when trained on data similar to their targets, their efficacy may be substantially reduced when applied to more diverse targets.

In perspective, besides the consistency of *foldare-predict* performance, it should be mentioned that improvements (compared to standalone), are generalizable when our two-step strategy is applied. For example, with RNAstructure: the standalone RNAstructure applied to the 25db dataset returned a consensus score of 0.697, a value that improved to 0.723 when our algorithm was run only on RNAstructure, i.e. when RNAstructure was used both as an ensembler and a predictor; to a lower extent, this was also true for other methods, as most combinations returned improved scores over the standalone predictions (**Figure 2**). Altogether, these results suggest that the information contained within structural ensembles (even when generated by the same prediction program) can be further leveraged for downstream refinement; for instance, through our pseudo-SHAPE co-input protocol, this ensemble-derived information can be reintroduced to improve the accuracy of final predictions, effectively enabling a second pass of structural information integration; to be noted, the efficacy of this second pass depends on model selection within the structural ensemble. Rather than using data from the complete Step 1 ensemble, the pseudo-SHAPE co-input is derived by parsing only the top *N* scoring models (*N* = 75 for foldare-predict, as empirically determined in Fig. 2B).

While a definitive theoretical explanation remains to be fully elucidated, we hypothesize that this second pass of information—filtered through a high-confidence sub-ensemble—effectively guides the downstream prediction toward more stable, high-confidence structural positions. In more general terms, the algorithm presented in this work represents an early example of a two-step strategy in which machine learning is first used to generate data-driven structural constraints (e.g., pseudo-SHAPE reactivities), followed by thermodynamics-based refinement. This approach could be considered conceptually analogous to template-based modeling in protein structure prediction, where force-field simulations are applied to experimentally derived or homology-modeled templates to refine their conformation. This study, in this sense, can only be considered a first step, since machine learning was not specifically trained for this task (rather, we used already existing, and pre-trained methods and models). However, our strategy could indicate a new line of development where machine learning ensembles, trained *ad hoc* (models for different cases, e.g. a model based only on data for human mRNAs, from icSHAPE in human cell lines), are used to guide downstream thermodynamic-based assessments, to create a model that is both probabilistically favoured and energetically sound.

Advances in tertiary structure analysis, coupled with efforts to broaden the experimental representation of RNA molecules in structural databases, are crucial for uncovering the details of RNA structural architecture — ultimately enabling the integration of all structural elements, such as stems, loops, and pseudoknots, into complete 3D models (Bu et al. 2025; Kwon 2025).

At the same time, we anticipate sustained interest in improving standard secondary structure prediction, particularly for high-throughput omics applications. These settings typically involve large-scale sequence datasets and require rapid, yet informative, structural overviews. For instance, many studies aim to quantify double-stranded RNA (dsRNA) content across samples (e.g., patient vs. control), as elevated dsRNA levels can trigger immunological responses. In such cases, a broad assessment of secondary structure distributions across RNA populations is often sufficient and still highly informative. With this work, we aim to contribute to this need by enhancing the accuracy and utility of ensemble-based secondary structure predictions.

Specifically for this task, we developed FoldARE, a computational tool that is *(i)* an implementation of our novel predictive algorithm (*foldare_predict*) and *(ii)* an analytical instrument for comprehensive structural investigations, employing the four established external methods considered in this study, in any user-defined combination. Regarding its predictive capabilities, *foldare_predict* runs by default with the optimized parameterization described in the results section; however, any predictor-ensembler combination can be explored, alongside variations in key parameters (such as ensemble size, thresholds, and weights) enabling completely customizable two-step strategy searches and replication of the analyses shown in Fig. 2.

Among its analytical capabilities, *FoldARE* allows cross-tool comparisons (to assess the similarity and divergence of predicted methods) and consensus-based rankings that aggregate results from different methods, which can be useful for “grading” the reliability of predictions (based on the extent of consensus, local or global); such comparative analyses at the ensemble level represent a significant addition to the current landscape of solutions for RNA secondary structure evaluations, particularly as *FoldARE* is available both as a standalone tool (for command line usage, e.g. for high-throughput) and as a user-friendly web application.

Then, a specific mention is due to the analysis of RNA modifications, in particular m6A; with a dedicated “compare m6A” module, *FoldARE* takes in an RNA sequence and, given a list of modification sites, it automatically generates (modified and unmodified) structural models, reporting the effects triggered by m6A, at the level of best model (MFE) or at the ensemble level. It is important to note that ensemble analyses can be especially indicative to detect smaller changes that may not emerge with the minimum free energy structure, but can be traced as perturbations in the structural ensemble, e.g. m6A modifications triggering a re-distribution of alternative models, favoring conformers with increased levels of single-strandedness. This fully automated workflow generates intuitive, interactive graphical outputs, enabling users – even those with little or no computational expertise – to thoroughly compare the effects of m6A modifications using one or both of the currently available methods (ViennaRNA, or RNAstructure), in the latter case enhancing reliability through cross-validation. While this is an accessory feature of *FoldARE*, it is, to our knowledge, the first implementation of this kind, with potentially impactful benefits to the ever-growing field of RNA modifications.

## METHODS

### Predictors and Ensemblers

The methods for our work were selected on the basis of some necessary features, theoretical and practical: *(i)* output to include detailed ensemble information, expliciting alternative conformations in standard dot bracket notations (i.e. pseudoknot-free), *(ii)* ability to utilize SHAPE-formatted co-input for guided predictions, *(iii)* capable of working with any input size (an asset for omics purposes, e.g. including lncRNAs or mRNAs), and *(iv)* free to download, use (customizing the search) as a command line program.Four methods were fitting these restraints: from the ViennaRNA package (v2.6.4), we used two programs, RNAfold, as a predictor, and RNAsubopt as an ensembler; they share the predictive approach, but RNAfold is optimized for a “best structure” output (Minimum Free Energy, MFE; Maximum Expected Accuracy, MEA, if chosen) and is capable of handling SHAPE-co input text files, while RNAsubopt is optimized for ensemble production (including specific command line customization). We used default parameters for RNAfold; for RNAsubopt, default parameterization was used except for the ensemble size (set to vary during optimization runs; and set to 75, as an optimized parameter) and for the ordering of alternative conformations; we tested both Zuker ordering (from most to least energetically favored) and Boltzmann sub-sampling (with *-N* option), specifying the number of non-redundant alternative conformations to extract, according to their probability from the partition function (i.e. these *N* unique structures represent the most statistically viable and lowest-energy alternatives). While our tools allow users to configure this choice, we selected Boltzmann sub-sampling for RNAsubopt in the main analysis (in Fig. 2) because it provided better overall performance, yielding slightly improved results with significantly faster computation (particularly with larger RNAs, e.g. > 1000 nucleotides).

From the RNAstructure package (v6.5), the Fold program was used with default parameterization, except for the ensemble size (set to vary, then set to 75); Fold returns an explicit ensemble file, where the first structure is the MFE (and the following structures are ordered by energy). LinearFold was used with the default -c option (for Contrafold) as a standalone predictor and as an ensembler; in the latter case, Zuker ordering was selected. Instead, when employed as a predictor LinearFold is run with the -v option (necessary to allow the employment of a SHAPE co-input text file). EternaFold was used with default parameters, except for the ensemble size (set as a variable, as in the other three programs). As representatives of deep learning methods, Mxfold2 and SPOT-RNA were chosen. Both are freely available as command-line tools and are well-maintained, with active links and straightforward installation following the authors’ instructions on standard Linux distributions (tested here on Ubuntu 22.04 LTS and 24.04 LTS). Additionally, both programs can accommodate large inputs, although SPOT-RNA is significantly more resource-intensive than the other methods evaluated in this study when processing larger RNAs. Following installation, both tools were executed using their default parameterizations.

The pseudo-SHAPE co-input was prepared internally by our method, as explained below.

### Preparation of the pseudo-SHAPE co-input

The key idea in this work is the use of in silico-generated pseudo-SHAPE co-input files. All models in a structural ensemble are parsed position per position, to count for single-strand RNA (ssRNA) occurrences, then transformed into a frequency, f(ssRNA); for example, for position *i*, with 10 models, half of which present a single strand prediction, f(ssRNA)*_i_* = 0.5. In our search, we employed a parametric scheme, with three cut-offs for high, higher-medium, and lower-medium values of single strandedness, termed x1, x2, x3, respectively. Given a position *i*, its frequency value, f(ssRNA)*_i_*, is assessed: depending on where it falls, with respect to x1, x2, x3, three multipliers are applied, K1, K2, K3, respectively. For example, if x2 < f(ssRNA)*_i_* < x1, a score equal to x2 * K2 is assigned to the position and written in the pseudo-SHAPE co-input; to further exemplify with numerical values: given the cut-offs set to x1=0.9, x2=0.5, x3=0.3, and the multipliers set to K1=2, K2=1.25, K3=0.75, a f(ssRNA)*_i_* = 0.6, positions in the intermediate bin (x1> 0.6 > x2), and thus it is evaluated by applying the K2 (K2=1.25) multiplier, for a pseudo-SHAPE value of 0.75 (1.25 * 0.6), which is then written out in the pseudo-SHAPE co-input. If instead, f(ssRNA)*_i_* = 0.4, which falls in the range of x3, then the multiplier is K3, and a value of (K1* f(ssRNA)*_i_*) 0.75 * 0.4 = 0.3 is written out. If a position has a f(ssRNA) < x3, then no value is added to the pseudo-SHAPE co-input. For a visual representation, we refer to the Results section (**Figure 1** and related discussion).

We run two optimizations; in the first varying the parameters with the following scheme: x1={0.9, 0.8,0.7}, x2={0.6,0.5}, x3={0.4, 0.3, 0.2}, K1={3, 2.5, 2}, K2={1.25, 1}, K3={ 0.5, 0.75}, ensemble size = {50, 75}; in the second, we refined the search by using the following scheme: x1 ={0.8, 0.85, 0.9}, x2 ={0.5, 0.55, 0.6}, x3 ={0.3, 0.35, 0.4}, K1 ={1.75, 2, 2.25}, K2 = {1, 1.25}, K3 = {0.5, 0.75}, with ensemble size =75. The pseudo-SHAPE guide co-inputs are written out in a two-column text format: each line contains the nucleotide index (*i*) and its pseudo-SHAPE value; this format is directly accepted by RNAstructure, ViennaRNA, and LinearFold; for EternaFold, the text files need to be converted to the BPSEQ format (with the same information).

### Scoring system

We employed and tested three different scoring systems: the first, consensus scoring A (scoreA), is an identity score; it returns the % of identical prediction:

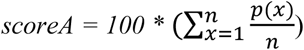

where n=number of positions, and p(x)=1, if the x-th position is conserved; else, p(x)=0. This score (consensus score, used as default) allows for fastest calculations while maintaining discriminatory power (Fig. 3D, and related discussion): it was designed to better suit large scale analyses (such as comparative omics, particularly when including long RNAs).

The second, scoreC, is a similarity score; it considers a window of 5 nucleotides, centered around the *x-th* position: if identical, counted as +1, else, proportionally contributing (e.g. if 3 out of 5 positions are identical, counted as +0.6); moreover, to account for possible small and local “shifts” we set a sliding window of -1 and +1: so, for each position, x, three 5 nucleotides windows (centered around *x*-1, *x*, *x*+1 respectively) are assessed for similarity with the reference structure: the best scoring window is taken (normally the one centered around *i*), and counted accordingly (i.e. in the example above, +0.6).

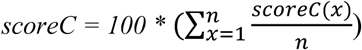

where n=number of positions, and

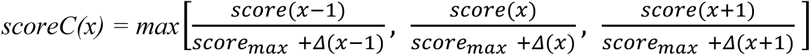

with score (x) = number of identities in the 5-nt window (x-2, .X+2), score_max_ = max value for score(x), with score_max_ = 5, given a window size of 5 (as employed here), and Δ(x) = score_max_ – score(x).

Finally, a matching bracket scoring scheme (scoreD) was considered: it represents a stricter version of an identity scoring, as it considers not only that brackets are predicted correctly, but also that they pair with the same partner in the sequence. This scheme, differently from the others, allows to define an F1 score (as here you can define false positives); however, it is less flexible in evaluating overall structural similarities (e.g. cases with a single variation, skew the pairing penalizing any downstream similarity). It works by listing all predicted pairs (e.g. [3, 24], pairing of the 3^rd^ position with the 24^th^) in the reference structure (ref_pairs), and in the predicted structure (pred_pairs). Then it accounts for pairs in common, as true positives, TP= pred_pairs & ref_pairs; and for pairs only in prediction (not in reference), as false positives, FP = pred_pairs - ref_pairs; and pairs only in reference as false negatives, FN = ref_pairs - pred_pairs. From this, we evaluated a scoreD, exactly as an F1 score:

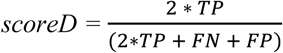

with TP = True Positives, FN = False Negatives, FP = False Positives. It should be noted that while the default scheme in FoldARE is scoreA, the other scoring systems are also available in the standalone version and can be set as default via the main configuration file (for any comparative analysis, e.g., for the aggregate score ranking in compare_all).”

### Other datasets

For testing and benchmarking we employed different datasets. For testing, we employed a manually curated non-redundant dataset of 25 RNAs, derived from RNAstrand (Andronescu et al. 2008) (15 RNAs, that had to comply with: 1500 nt > length > 50 nt, and at least 20% of dsRNA) and 10 additional, more recent and manually selected, structures from the Protein Data Bank. Entries in this dataset (25db; see Table 1 for a complete list) are characterized by a sequence identity < 60% between any of the models, with an average length of 354 nt (13 RNAs> 200 nt, and 5 < 100 nt). In building this testing dataset, we aimed to capture as much diversity as possible (most RNA structural databases are excessively redundant for the same types of RNA, as described in the Introduction), while keeping the dataset manageable for computationally intensive optimization runs. Larger datasets included non-redundant (at 80% sequence identity) models, with standard dot bracket notation (no pseudoknots, at least 20% dsRNA content) from: RNAstrand (n=424 models), Archivell (n=825) (Sloma & Mathews, 2016), bpRNA (n=715) (Danaee et al. 2018). It has to be noted that, while for the others we took the whole dataset, for the bpRNA case, being much larger (> 20,000 entries), we randomly selected 1,000 RNAs, then subjected to the filtering described above; this procedure was chosen in order to have similar numerosity among different datasets (and thus avoid skewing the analysis toward the more abundant dataset). Finally, we included a recently published dataset, eFold, available as a github (https://github.com/rouskinlab/eFold) from the Rouskin lab: this is currently the only dataset available of this type (with secondary structure annotation, as standard dot bracket) for human data, including mRNAs; here we employed 500 randomly selected (out of 1455) mRNA entries and pri-miRNA (out of 1098), and all the lncRNA (n=24) datasets. Complete details on the models employed in our analyses are available as Supporting information (Table S1, S3).

The performances against these datasets are reported in **Figure 4**, as Z-scores; this evaluation is useful to homogeneously compare data across different datasets: the Z-scores are calculated as follows:

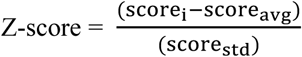

Where score_AVG_ is the average of the scores of all methods, and score_STD_ is the standard deviation of the scores of all methods, so that positive Z-score values indicate better than average methods, while negative Z-score values, below average methods.

## Contributions

SMM: Conceptualization, Methodology, Software, Data curation, Validation, Writing – original draft; VH: Software, Data curation, Validation, Writing – review & editing; TT: Resources, Supervision, Writing – review & editing

## Acknowledgments

We thank Stephanie Halene and Christian Ramirez for their valuable comments and discussions, Riccardo Scandino and Valter Cavecchia for the support with the web server. This study was funded by AIRC under MFAG 2020 (ID. 24883 project), the MUR PNRR project CN RNA&GT RINGTAIL (CN00000041) M4C2 Inv 1.4 funded by the NextGenerationEU, Fondazione VRT (“bando intelligenza artificiale 2024”). This work was also supported by the “Departments of Excellence 2023-2027” initiative (Law 232/2016), project no. 40613, funded by the Italian Ministry of University and Research (MUR). We are also grateful to AIL Trento and AIL Bolzano for their generous financial support.

## Notes

https://github.com/TebaldiLab/FoldARE/

https://rdds.it/foldare/

### Competing Interest Statement

The authors have declared no competing interest.

### Summary of Updates

The FoldARE web server is now available.

